# Collecting large datasets of rotational electron diffraction with ParallEM and SerialEM

**DOI:** 10.1101/2020.06.01.128793

**Authors:** Kiyofumi Takaba, Saori Maki-Yonekura, Koji Yonekura

## Abstract

A semi-automated protocol has been developed for rotational data collection of electron diffraction patterns by combined use of SerialEM and ParallEM, where SerialEM is used for positioning of sample crystals and ParallEM for rotational data collection. ParallEM calls standard camera control software through an AutoIt script, which adapts to software operational changes and to new GUI programs guiding other cameras. Development included periodic flashing and pausing of data collection during overnight or day-long recording with a cold field-emission beam. The protocol proved to be efficient and accurate in data collection of large-scale rotational series from two JEOL electron microscopes, a general-purpose JEM-2100 and a high-end CRYO ARM 300. Efficiency resulted from simpler steps and task specialization. It is possible to collect 12–20 rotational series from ∼ −68º to ∼ 68º at a rotation speed of 1º /s in one hour without human supervision.

## 1. Introduction

Automation in data collection is becoming more and more important in many scientific fields. This trend is being seen in both cryo-electron microscopy (cryo-EM) and X-ray crystallography. Indeed, automated image-taking is essential technology for high-resolution single particle cryo-EM, as it requires several tens of thousands to a million molecular images selected from thousands of image stacks. Computer programs such as SerialEM (Mastronarde, 2005), Leginon (Suloway et al., 2005), EPU (Thermo Fisher), JADAS (JEOL) and Latitude S (GATAN) have been developed for this purpose, and are widely used in laboratories and shared facilities. A typical protocol comprises roughly three steps: calibration and alignment of the microscope; selection of suitable areas of grid squares and often carbon holes; and automatic data collection. Macromolecular X-ray crystallography (MX) has also benefited from automation in data collection, and merging of many datasets has revealed complex protein structures, which would otherwise be difficult to solve (e.g. Huang et al., 2018; Hori et al., 2018).

A similar approach could apply to electron 3D crystallography (also known as 3D ED / MicroED). In a standard way of this technique, sequential frames of rotational diffraction patterns are recorded on a detector using an electron microscope, while the sample crystal is continuously rotated (e.g. Nannenga et al., 2014; Yonekura et al., 2015; Clabbers et al., 2017) as in MX. In automated MX data collection, small wedges of a rotational series are collected from many small crystals or parts of larger crystals to reduce radiation damage to the sample. Indexing of diffraction spots is usually possible from these datasets, as different intersections of the 3D Fourier space in the direction of the incident X-ray beam are recorded in a single frame on the detector at a wavelength of ∼ 1 Å due to the curved surface of the Ewald sphere. In contrast, reflecting the much shorter wavelength of electrons (∼ 1/50 of the X-ray wavelength), electron diffraction yields an almost flat Ewald sphere, resulting in each frame having only a single intersection or limited intersections normal to the incident beam. Thus, electron diffraction patterns should be collected from relatively larger wedges except for samples with known lattice parameters (Smeets et al., 2018; Bücker et al., 2020). These large datasets could be processed and sorted with programs such as KAMO (Yamashita et al., 2018) originally developed for MX, and Instamatic / REDp (Cichocka et al., 2019; Wang et al., 2019) specially for electron diffraction.

There have been several reports and programs for automated data collection of rotational electron diffraction patterns (e.g. Smeets et al., 2018; Cichocka et al., 2019; Wang et al., 2019; de la Cruz et al. 2019; Bücker et al., 2020). De la Cruz et al. (2019) proposed the use of SerialEM, which has become one of the most popular programs for automated data collection in the cryo-EM field and is available with the JEOL, Thermo Fisher Scientific and HITACHI electron microscopes. Although a script program of SerialEM was provided with the report (de la Cruz et al., 2019), its use requires a specific socket connection to the Thermo Fisher Scientific electron microscope and is thereby limited to Thermo Fisher Scientific machines (de la Cruz et al., 2019).

We have developed an alternative solution by combining SerialEM and ParallEM, the latter being a GUI program package for controlling and monitoring the JEOL electron microscope (Yonekura et al., 2019; Hamaguchi et al., 2019). Our approach fulfills the required functions described above with increased usability and high customizability partly thanks to task specialization and incorporation of the standard camera control software through the GUI operation scripting language AutoIt (https://www.autoitscript.com/site/). We here report details of this approach and its development and provide a working flow scheme that facilitates data collection of rotational electron diffraction patterns. These programs and scripts can be used for general-purpose and high-end microscopes, and also supports tasks associated with flashing for overnight or day-long data collection with a cold field-emission beam.

## 2. Setup and Development

### 2.1 Hardware setup

In this study, rotational diffraction patterns were collected using a JEOL CRYO ARM 300 electron microscope on a bottom-mounted scintillator-based CMOS camera, TVIPS XF416, and using a JEOL JEM-2100 microscope mounted on a Gatan OneView IS camera, which has also scintillator on its CMOS sensor. The CRYO ARM 300 microscope is equipped with a cold-field emission gun operated at an accelerating voltage of 300 kV, quad condenser lenses for parallel illumination and an in-column Ω-type energy filter. The JEM-2100 microscope has a LaB6 gun operated at 200 kV accelerating voltage, and diffraction patterns were taken with a GATAN Elsa cryo-transfer sample holder on this machine. Most JEOL microscopes should be available with the TEM External library (see below) for the data collection method developed and proposed in this manuscript.

### 2.2 Software setup

The proposed method uses ParallEM (Yonekura et al., 2019; Hamaguchi et al., 2019), SerialEM (Mastronarde, 2005), AutoIt (https://www.autoitscript.com/site/) and camera control software such as DigitalMicrograph (Gatan), and EMMenu (TVIPS). SerialEM Version 3.8.0beta8, AutoIt V3, DigitalMicrograph GMS 3 (3.21.1365.1) and EMMenu 4 (5.0.12.0) were tested on Microsoft Windows 7, 10 and Server R2.

We have developed the ParallEM package (Fig. 1; Yonekura et al., 2019; Hamaguchi et al., 2019) that consists of GUI programs for controlling and monitoring the JEOL electron microscope through the JEOL TEM External library. The TEM External library is a collection of basic function calls to change the setting of the JEOL microscope and fetch the status from an external computer. The library is based on Microsoft .NET Framework, and we used TemExt3.0 2.1.6. SetDiff in ParallEM switches the lens condition (beam spot size, illumination angle α and brightness, diffraction focus, and beam position) between either search for crystals or electron diffraction data collection (Fig. 1A). Rotation in the package adjusts the speed of the rotation and synchronizes open / close of the beam blanking shutter with tilt angles at the start / end points (Fig. 1B). For reading the goniometer tilt angle, Rotation automatically switches to a rotary encoder, which continuously detects the goniometer angle, from the factory default potentiometer in JEOL microscopes, because the rotary encoder provides a more accurate angle than the potentiometer. In addition it has a function that saves the rotation range, hexadecimal values of the lenses and deflector coils and related information including a used condenser aperture, selected-area aperture and energy slit in / out for CRYO ARM 300 in a folder “ParallEM¥Rotation” at the desktop on start of taking each rotational series. ChkLensDef checks if the settings of lens and deflector coils are correct (Fig. 1C). Flashing controls the flashing and counts the time (Fig. 1D). SamplePositioner records the trajectory of the stage movement in diffraction data collection. Default values of lens, deflector coils, and rotation degree / pulse of the goniometer may differ from microscope to microscope, and machine-specific settings can be fed from configuration text files for ChkLensDef and Rotation. The ParallEM programs are all written in C#.

**Figure 1.**
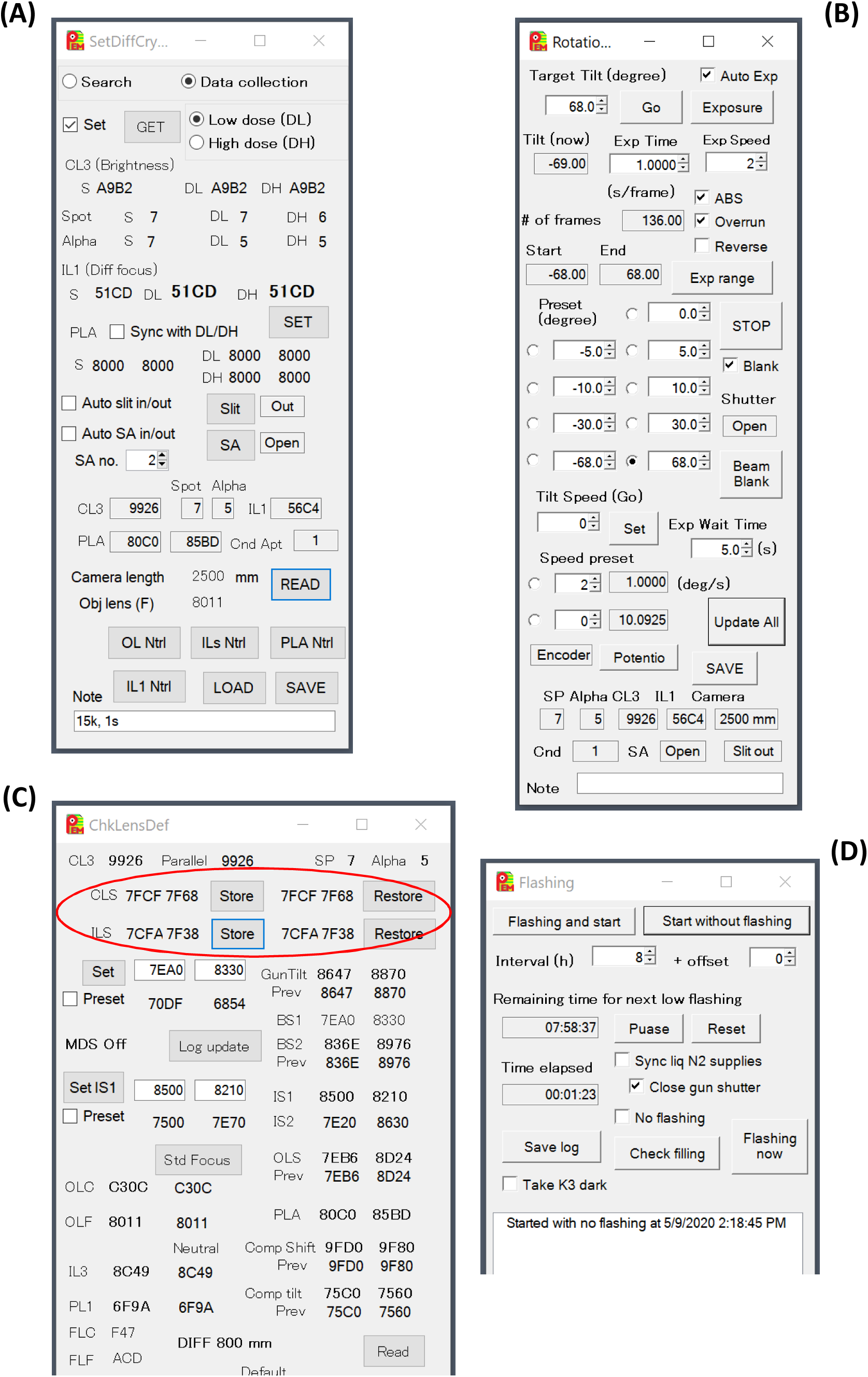
GUI programs available for data collection in electron 3D crystallography (included in the ParallEM package). (A) SetDiff. Selection of the radio buttons “Search” and “Data collection” switches lens conditions among a search mode (defocused diffraction mode) and two data-taking modes (focused diffraction modes) that store two different lens settings usually at lower and higher doses, referring to the “Low dose (DL)” and “High dose (DH)” radio buttons, respectively. A push on the “GET” / “SET” button fetches / sets the lens conditions from / to the microscope. If the box “Set” is checked, the lens condition is switched upon change in selection of the “Search” or “Data collection” radio button. (B) Rotation. Pushing the “Exposure” button starts gonio rotation after waiting for the time specified in the “Exp Wait Time” text box and synchronizes shutter opening / blanking with tilt angles at the “Start” / “End” setting angles. The rotation speed is set to the value in “Exp Speed". A check in the “Auto Exp” box calls batch job “RotationAutoExp.bat”, which then calls an AutoIt script such as “MovieOneView.au3” and “MovieEMMENU.au3” to start movie recording with DigitalMicrograph and EMMenu, respectively. A push on the “Go” button rotates the goniometer to the value in “Target Tilt” at a rotation speed in “Tilt Speed (Go)” without referring to the shutter. (C) ChkLensDef. Red circle encloses the functions to store / restore x, y values of “CLS” and “ILS” for each beam spot size, α, and camera distance (only with ILS) implemented in this study. See Fig. 3 and Yonekura et al., 2019 for SamplePositioner and see also Yonekura et al., 2019 and Hamaguchi et al., 2019 for more details on ParallEM. (D) Flashing. It carries out flashing in a given period and replies with the remaining time till next flashing when there is a call from a batch file.

The data collection method developed here combines SerialEM for positioning of sample crystals and ParallEM for rotational data collection. SerialEM is a widely used program for automated data collection of electron tomography and single particle analysis. The application can also be used for rotational data collection of diffraction patterns with Thermo Fisher Scientific microscopes (de la Cruz et al., 2019). However, some functions in the current version of SerialEM do not work well with the JEOL microscope, and ParallEM can complement these. Also, due to its multi-functionality, the usual setup of SerialEM is a time-consuming task requiring knowledge and experience. In our application, a setup only requires the specific camera setting and stage calibration at low magnifications for taking a whole sample grid map.

AutoIt is a scripting language for GUI operations in Windows application programs such as mouse clicking and editing values in text boxes (https://www.autoitscript.com/site/). Any GUI elements such as button, text box, and slide bar on Windows programs have unique control IDs, and AutoIt uses the ID to operate the GUI programs. In our application, AutoIt sends actions on GUIs to the camera control software from Rotation in ParallEM (Fig. 2).

**Figure 2.**
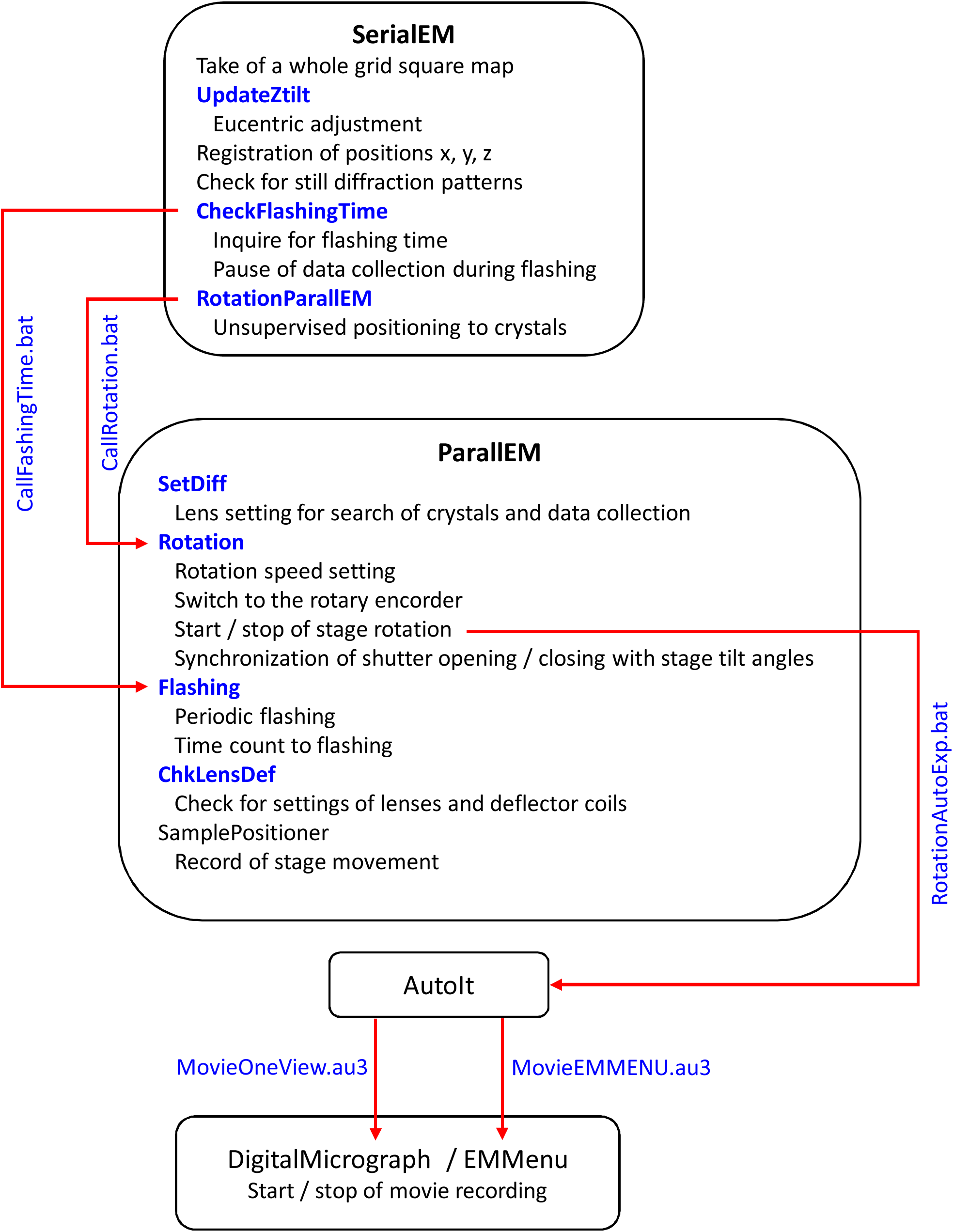
Schematic diagram of the approach combining ParallEM and SerialEM, developed here. Developments in this study are shown in blue.

### 2.3 Software Development

We have implemented functions for adoption of automated data collection in the ParallEM programs, Rotation and Flashing. The functions can be carried out from SeriallEM through a Windows batch file (see Supplementary Fig. S1). Thereby, Rotation takes user-specified parameters (exposure time / frame, target tilt angle, rotation speed and waiting time before tilt start) and launches an AutoIt script to start movie recording through GUI operations on the camera control software such as DigitalMicrograph and EMMenu.

We made AutoIt scripts for movie recording on the OneView IS camera with DigitalMicrograph and on the XF416 camera with EMMenu. The scripts MovieOneView.au3 and MovieEMMENU.au3 are provided in Supplementary Fig. S2. Control IDs of GUI buttons and text boxes may be different from ours depending on the user environment, but ID values can be easily checked with an accompanying tool called AutoIt Window Information Tool. It can be tuned without difficulty for any changes in operation of the camera control software and for adoption of new GUI programs for operating other cameras.

In the CRYO ARM 300 microscope, a cold field emission gun needs periodic flashing to refresh the tip of the gun and this obviates the substantial decrease in beam intensity. We set the interval between flashing cycles to 8 h (Hamaguchi et al., 2019) and found that the electron beam generated from the cold-field emission gun does not decay much during data collection of smaller-scale datasets. However, longer data collection requires flashing. SerialEM carries out flashing during automated data collection, but we sometimes stopped data collection and initiated the programs for sample exchange or due to unrecoverable errors in SerialEM and the camera control software, which reset the time for flashing. Flashing in ParallEM is a stand-alone program, and is set to keep running without interruption and displaying time counts to the next flashing. We implemented a function in Flashing and made a SerialEM script CheckFlashingTime (Supplementary Fig. S3B) to monitor the time remaining until the next flashing period and to pause data collection until flashing is over.

In order to incorporate the setup and development described above, we prepared a SerialEM script RotationParallEM (Supplementary Fig. S3C). Thanks to task specialization among ParallEM and SerialEM, the script and preparation before executing the script are simpler than those of CRmov (de la Cruz et al., 2019), which carries out rotational data collection with the Thermo Fisher ScientificI electron microscope. Our scheme could also significantly reduce operator labor through implementation of an automated eucentric adjustment script UpdateZtilt (Supplementary Fig. S3A). The concept of this protocol is illustrated in Fig. 2.

Apart from the development of semi-automated data collection as described above, we have improved the operationability of ParallEM. One of the improvements relates to the condenser lens stigmators (CLS) and the intermediate lens stigmators (ILS). Many lens conditions differ in diffraction and imaging modes in the CRYO ARM 300 microscope: We use the largest beam spot and larger illumination angles (weaker beam) for the former and small spots and small angles for the latter. Values of CLS change considerably depending on the spot size and illumination angle, but only single values can be kept in a particular setting. In diffraction mode, values of ILS change depending on the camera distance as well as the illumination angle. ChkLensDef can store / restore separate CLS and ILS values for all spot sizes and illumination angles (Fig. 1C). Separate ILS values are also kept for all camera distances. In addition, SetDiff has check boxes for automatic insertion of the energy slit and a selected-area aperture when switching to the data collection mode if the microscope supports those functions (Fig. 1A).

## 3 Working flow

This section describes our standard protocol for semi-automated data collection of rotational electron diffraction patterns using ParallEM and SerialEM. Our protocol consists of calibration and alignment of the microscope; selection of suitable crystals; and unsupervised data collection based on the selection. We do not use the “Low Dose Control” panel in SerialEM, as this mode in the current version appeared incapable of saving the intermediate lens 1 (diffraction focus) in JEOL microscopes and also, in the low dose mode, we often experienced unintended / accidental adjustments in lens setting probably due to misoperation.

1. Select “Open” in the Navigator menu of SerialEM and take an overview of the entire grid at magnifications 50 – 80 × from “Setup Full Montage” in the Navigator menu. It is convenient to make a copy of the entire map for reference from “New Window” in the Window menu.
2. Check on large ice contamination or broken carbon film as a landmark from “Add Points” in the Navigator dialog box. Go to the checked point from “Go To XY”.
3. Take an image covering one grid square that includes the land mark. We usually use 400 × for a 200 - 300 mesh grid. Keep this magnification until Step 7.
4. Left click on the landmark and apply a global shift between the entire map and the square map from “Shift to Marker” in the Navigator menu with keeping selection of the entry of the landmark.
5. Record the positions of good squares with crystals suitable for data collection in the entire map from “Add Points” in the Navigator dialog box, and set “Acquire (A)” to all the selected squares. It is easier to use the “Collapse” check box to treat the selected square items as a group.
6. Run the script UpdateZtilt from “Run script” in “Acquire at Items” in the Navigator menu to adjust the eucentric center for all the square maps.
7. Acquire all square maps for reference with “Acquire map image or montage” instead of “Run script” after marking each with “Acquire (A)” and checking “New file at group” in the Navigator dialog box. This step may be skipped if the entire map resolves crystals well.
8. Save Navigator items to a file.
9. Exit SerialEM for TVIPS cameras, as SerialEM and EMMenu interfere and cannot be run simultaneously. Go to Step 12 for Gatan cameras.
10. Launch a “No-Camera” version of SerialEM, where the TVIPS camera is removed from ActiveCameraList but a dummy camera entry is added in SerialEMproperties.txt. Thereby, SerialEM does not access the TVIPS camera, but allows control of stage movement in the electron microscope.
11. Select “Read & Open” in the Navigator menu to restore the Navigator items.
12. Set the lens condition to search mode with SetDiff.
13. Move to a marked square from “Go To XYZ”.
14. Search for a good crystal in live view with the camera control software or SerialEM.
15. Record the position from “Add Stage Pos” in the Navigator dialog box.
16. Repeat from Step 13 to register all positions of the crystals to be recorded into Navigator items. Mark all the positions in the Navigator dialog box with “Acquire (A)”.
17. Set the lens condition to data collection mode with SetDiff and make the camera control software ready for data collation.
18. Adjust beam stop to cover the center beam and insert a SA aperture and an energy slit if needed.
19. If needed, take still diffraction patterns for all the recorded points with SerialEM as in Step 7. To do so, exit EMMenu and the “No-Camera” version of SerialEM, and launch the normal setting of SerialEM.
20. Edit Navigation items based on the still patterns taken in Step 19.
21. Run the script RotationParallEM from “Acquire at Items”. The script repeats data collection for all the registered positions without human supervision, and stops after all the crystals are exposed and the rotational series saved in storage. The lens condition never changes during data collection.

Fig. 3 shows the display of data collection carried out in this manner with the JEM-2100 microscope and OneView IS.

**Figure 3.**
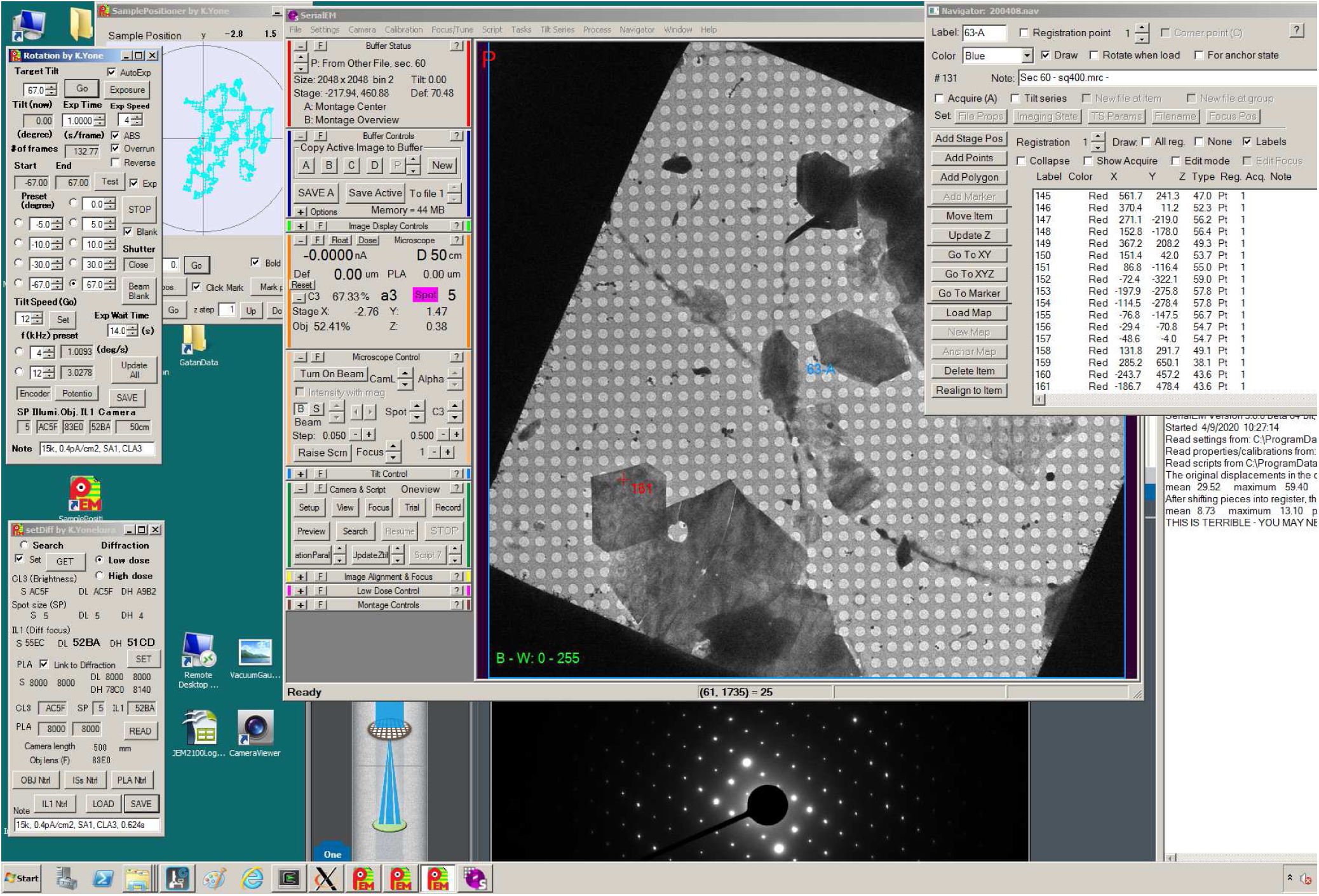
Display of data collection using the method developed in this study on the OneView IS camera with the JEM-2100 microscope. Rotational diffraction patterns were collected from thin crystals of an organic molecule.

## 4. Discussion

We have developed a semi-automated protocol for rotational data collection of electron diffraction patterns with JEOL microscopes by combining SerialEM and ParallEM, with AutoIt scripts engaging the default camera control software of ParallEM. The approach can be custom adjusted for changes in operation of the camera control software and adopts new GUI programs for operating other cameras. Thus, development of specific device drivers, which needs substantial time and labor and often introduces bugs, is not needed, and the operator can use the well-tested GUI-based software provided by the camera manufacturer.

Following schemes for single particle analysis (e.g. Mastronarde, 2005; Suloway et al., 2005) and in a previous report (de la Cruz et al., 2019), the developed approach consists of calibration and alignment of the microscope; selection of suitable areas; and unsupervised data collection based on the selection. The second part, which corresponds to registration of x, y, z positions of crystals to be exposed, is the most time-consuming. We carry out grid-square-based adjustment of z positions by a batch job of SerialEM using the auto eucentricity function (Step 6 in the working flow), and this adjustment is usually acceptable at the expected level of the current goniometer. Still, x, y positioning of crystals has to be done manually. We plan to develop a more sophisticated algorithm to reduce the amount of labor for this task. Switching among different observation modes never happens during data collection in the third part, and thus it is free from hysteresis in changes of lens and deflector coil values as in de la Cruz et al. (2019).

We tested the scheme on two JEOL electron microscopes, a general-purpose JEM-2100 and a high-end CRYO ARM 300, and found that data collection was highly effective and efficient on both machines. Even novice users were able to operate the programs thanks to their usability, and succeeded in obtaining good diffraction datasets. We usually collected one rotational series from ∼ −68º to ∼ 68º from a single crystal in ∼ 2 min 20 s at a rotation speed of 1º / s. Twelve – 20 rotational series / h can be collected with this setting. The approach rarely failed to locate the x, y, z positions registered for good crystals on both microscopes with an accuracy comparable to or better than full manual alignment for each crystal just before its data collection (Yonekura et al. 2019). A typical display of data collection is shown in Fig. 3 with rotational electron diffraction patterns from a thin crystal of an organic molecule. We processed and sorted these patterns with the automated data processing program KAMO (Yamashita et al., 2018) originally developed for processing and sorting many small-wedge datasets of X-ray diffraction patterns, and succeeded in solving structures beyond 1 Å resolution (manuscripts in preparation).

The CRYO ARM 300 microscope has a cold-field emission gun, which is not expected to have a significant effect on crystalline samples, but does require periodic flashing (Yonekura et al., 2019). Flashing in ParallEM and a SerialEM script CheckFlashingTime support periodic flashing and pausing of data collection during flashing for overnight or day-long data collection with a cold field-emission beam. The electron optics, including settings for parallel illumination, diffraction, and energy filtration in the CRYO ARM 300 microscope are stable over a single 1 - 3 day session, which consists of multiple flashing cycles and overnight breaks with the beam off.

In summary, the developed protocol is effective in accurate and efficient unsupervised data collection of rotational electron diffraction patterns from small and thin crystals of proteins and organic molecules. The ParallEM package including Windows batch files, SerialEM and AutoIt scripts is available on request. The script files are also available from a supplementary material of this manuscript.

## Acknowledgments

We thank Tasuku Hamaguchi for feedback on use of ParallEM, David Mastronarde for technical assistance with SerialEM, Radostin Danev for SerialEM setup in single particle data collection, Keitaro Yamashita for usage of KAMO, Satoru Inoue and Tatsuo Hasegawa for providing crystal samples of organic molecules. This work was partly supported by Japan Society for the Promotion of Science Grant-in-Aid for Scientific Research Grant 16H04757 (to K.Y.), Japan Society for the Promotion of Science Grant-in-Aid for Challenging Exploratory Research Grant 24657111 (to K.Y.), the RIKEN Pioneering Project, Dynamic Structural Biology (to K.T., S.M.-Y. and K.Y.), the Cyclic Innovation for Clinical Empowerment (CiCLE) from the Japan Agency for Medical Research and Development, AMED (to K.Y.), and JST CREST Grant Number JPMJCR18J2, Japan (to K.T., S. M.-Y., and K. Y.).

## Supplementary Information

**Supplementary Fig. S1.**
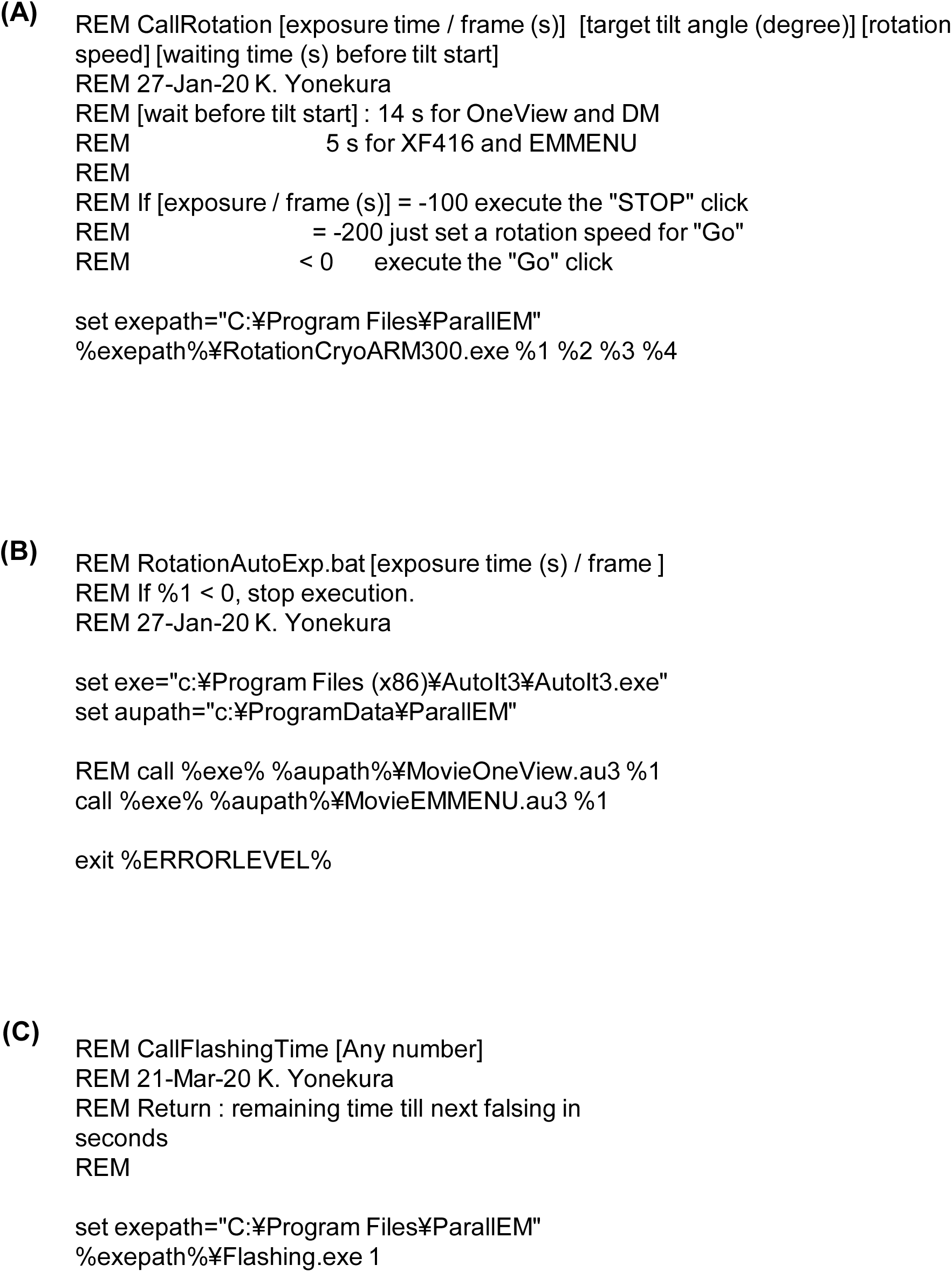
Windows batch files developed in this study. (A) CallRotation.bat. (B) RotationAutoExp.bat. (C) CallFlashingTime.bat.

**Supplementary Fig. S2.**
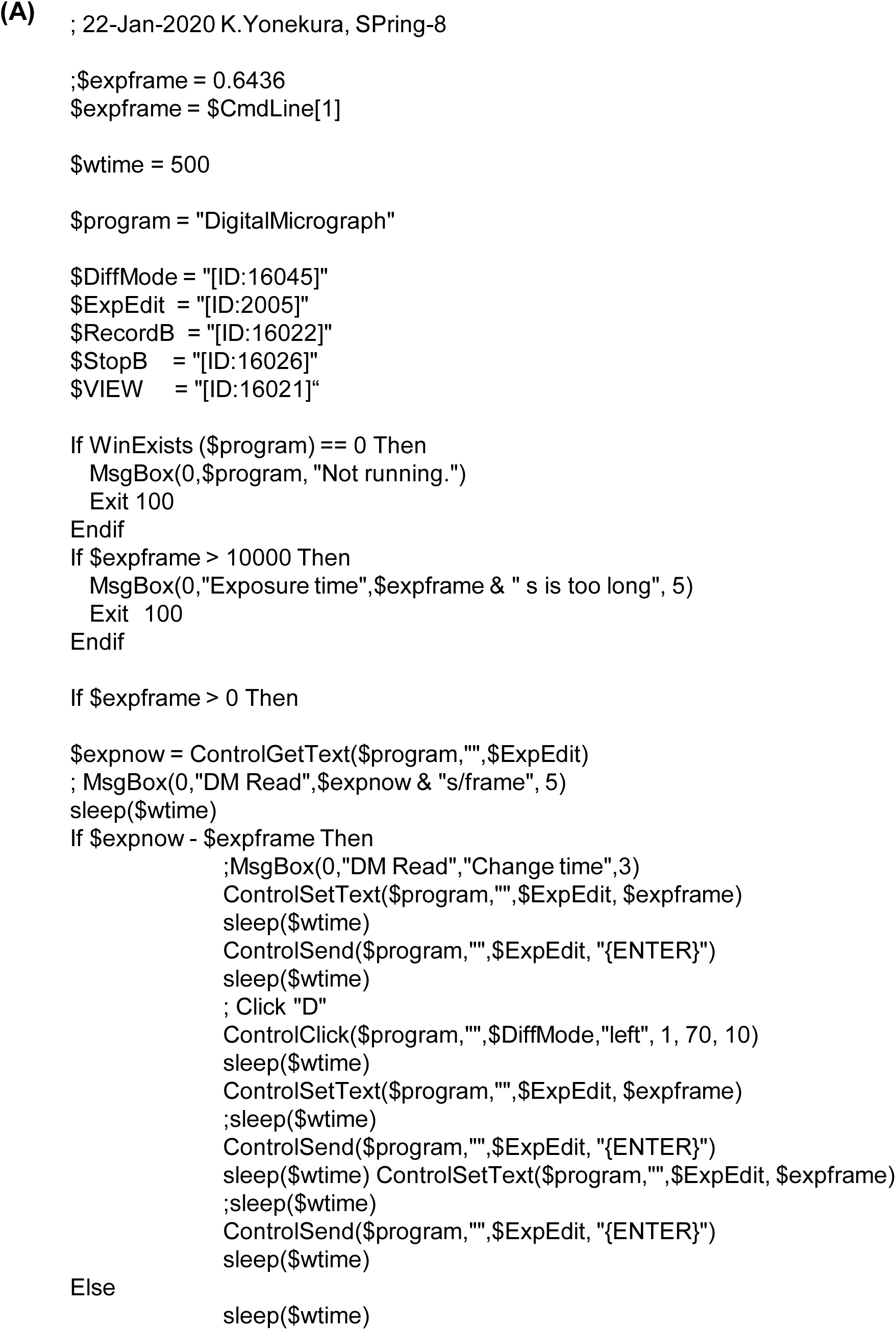
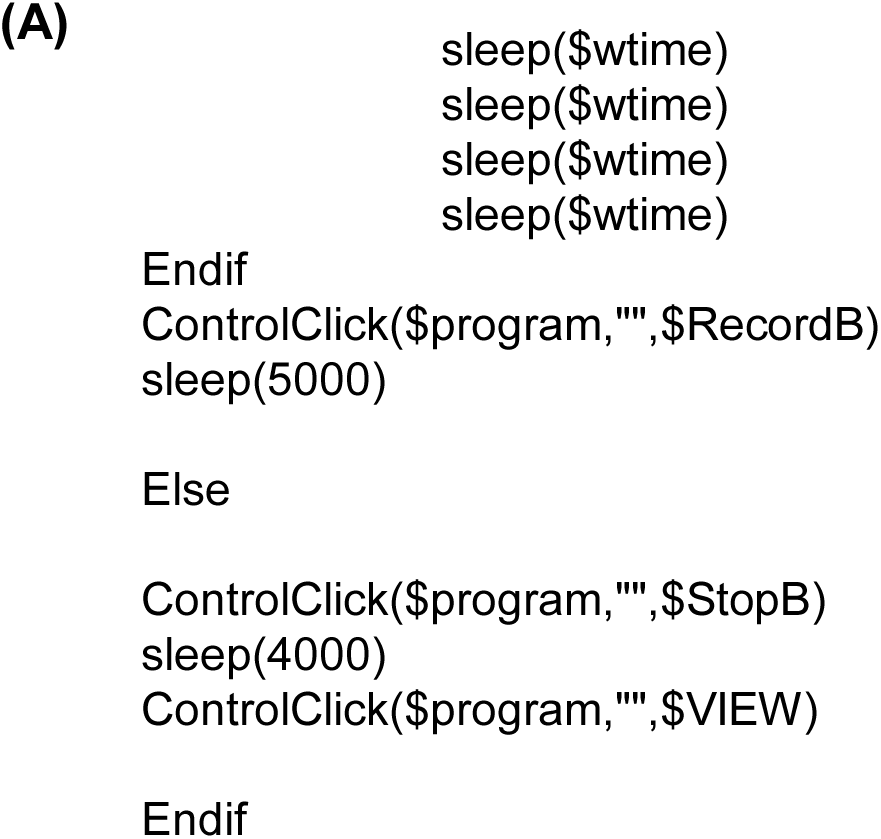
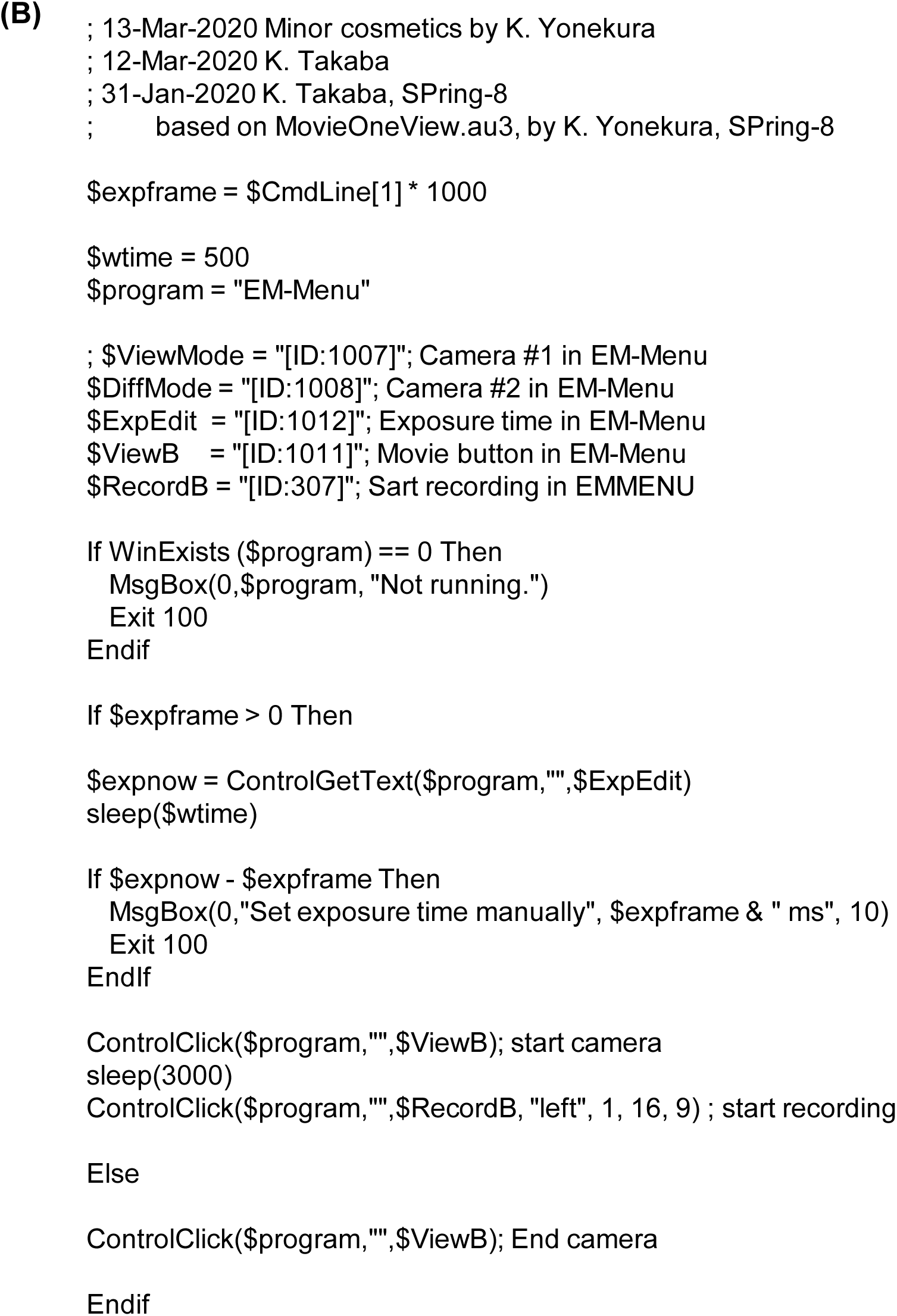
AutoIt scripts developed in this study. (A) MovieOneView.au3. (B) MovieEMMENU.au3.

**Supplementary Fig. S3.**
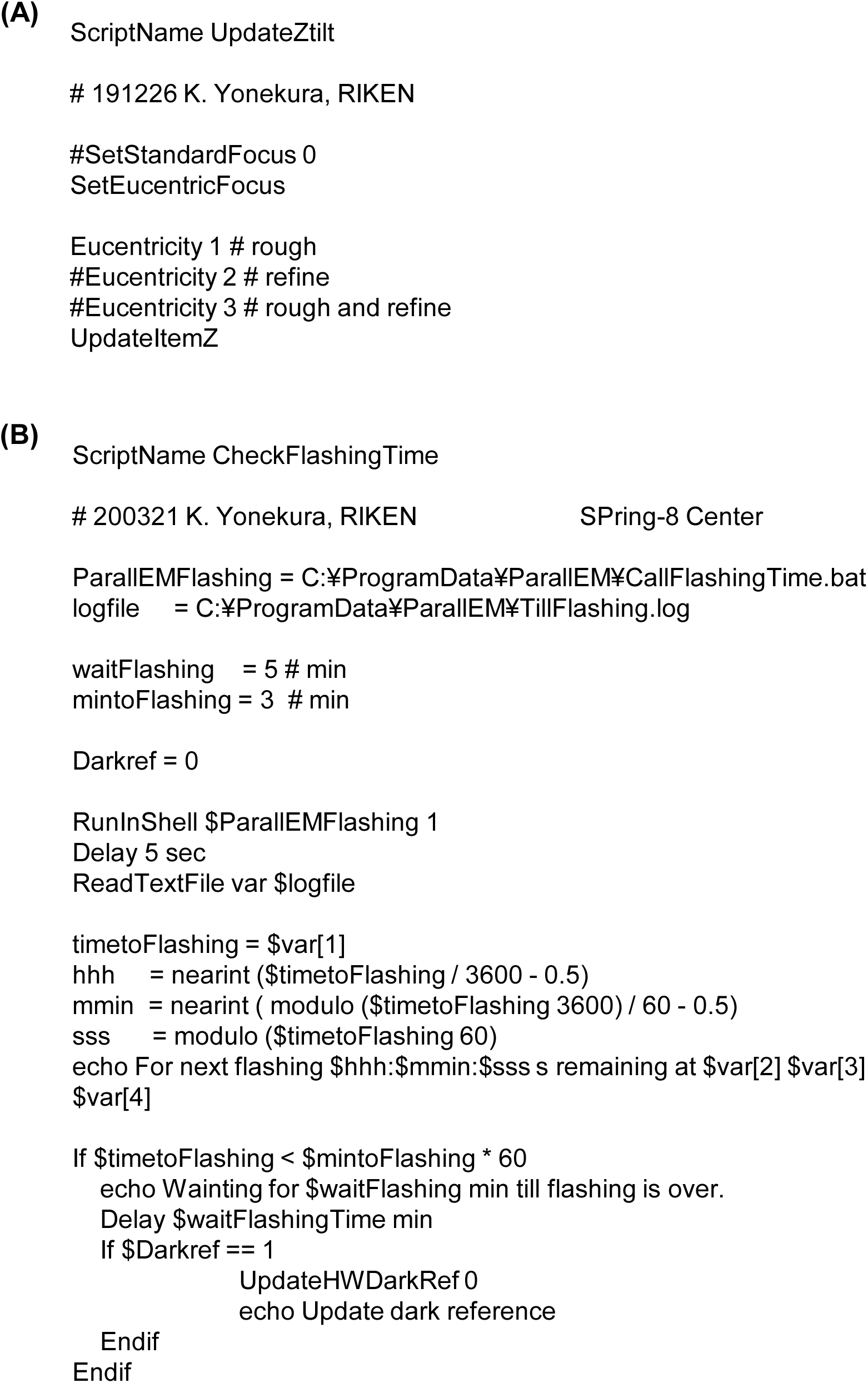
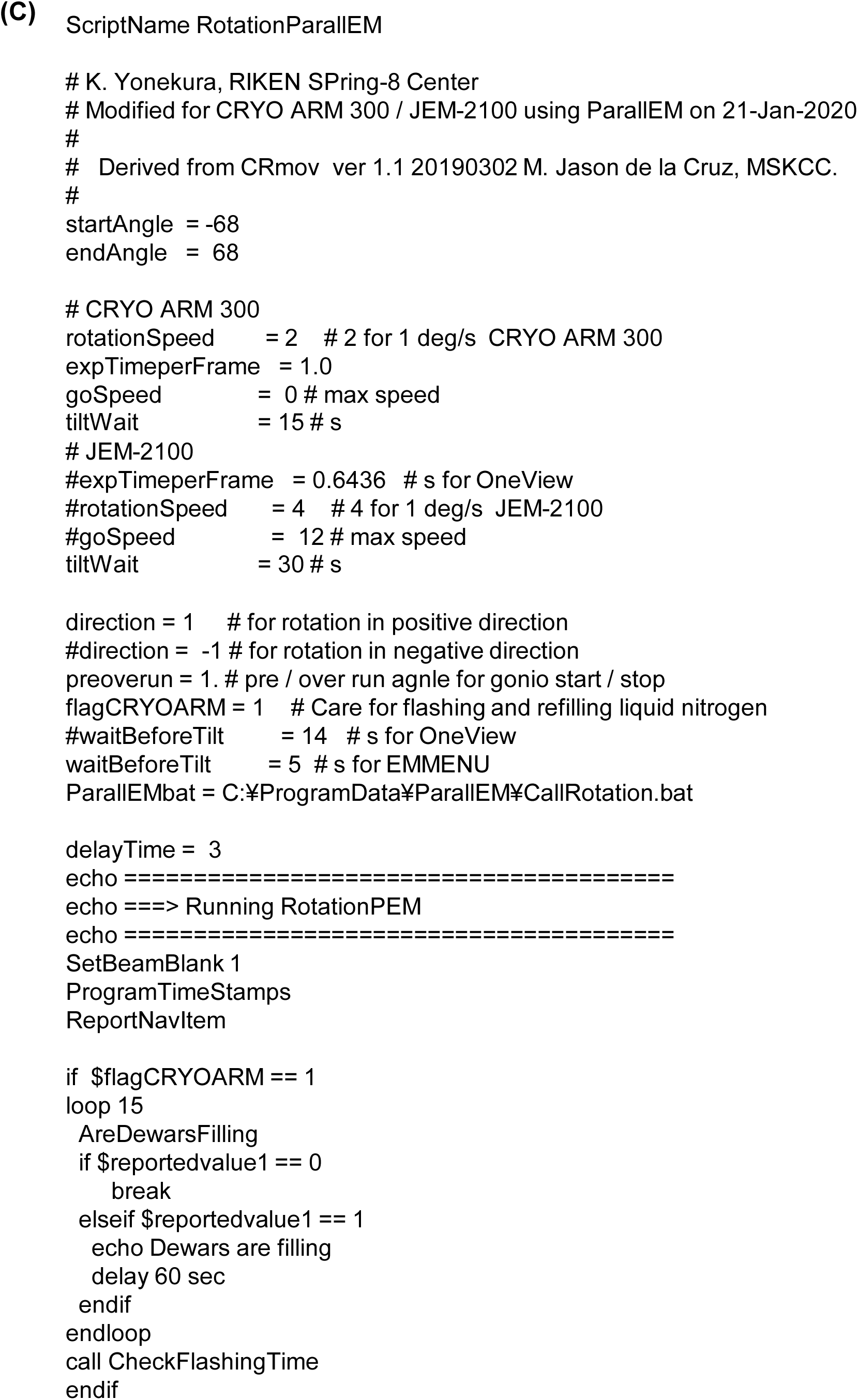
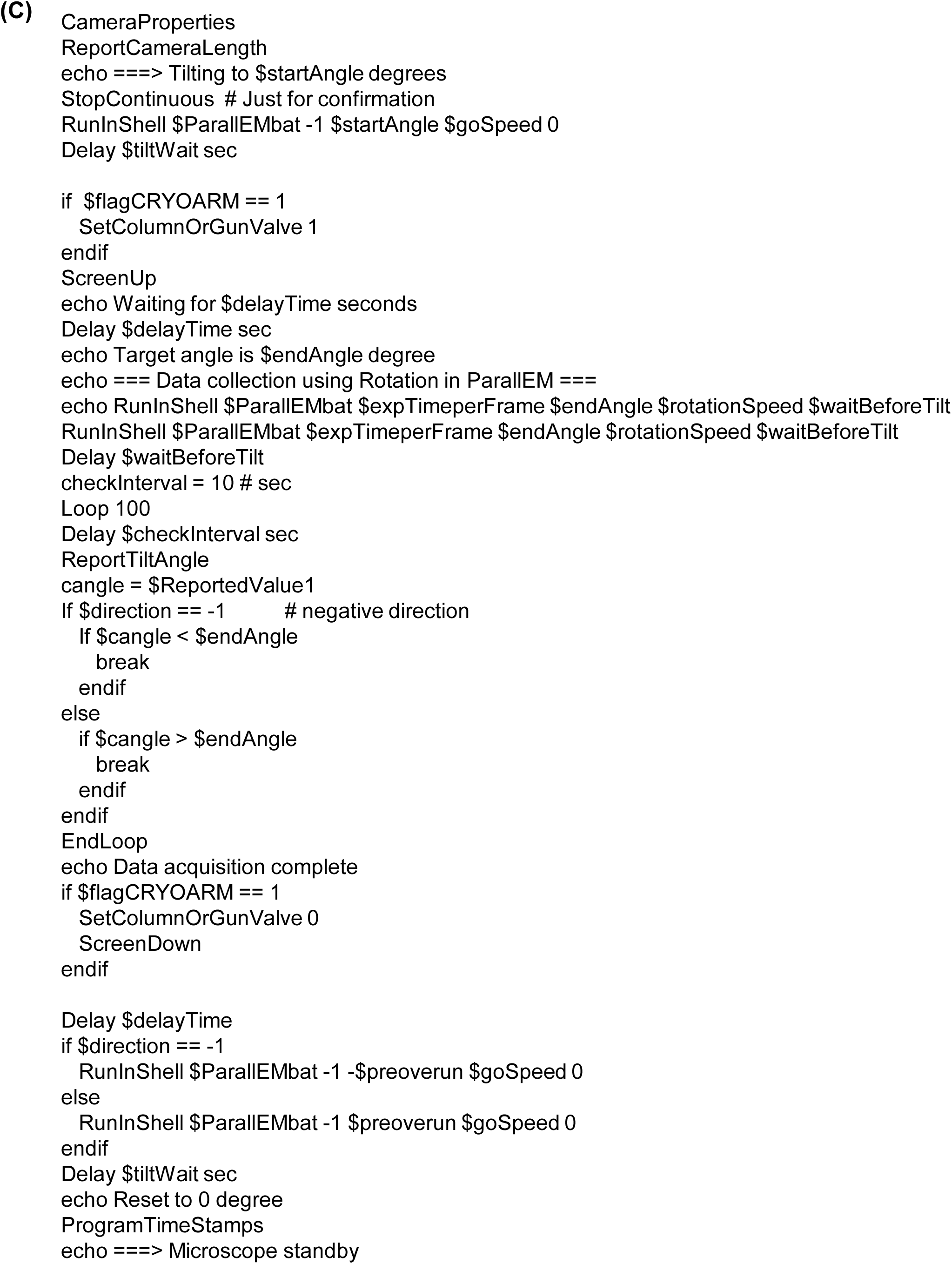
SerialEM scripts developed in this study. (A) UpdateZtilt.txt. (B) CheckFlashingTime.txt. (C) RotationParallEM.txt.

